# Root organogenesis induction in *Epipremnum aureum* stem cuttings with plant biostimulants and synthetic rooting hormone

**DOI:** 10.1101/2022.07.28.501829

**Authors:** D.E. Villafuerte, E. Angeles, A. Bayog, R. Duka, N.L. Meñoza, M.A. Sanchez, R. De Jesus

## Abstract

Plant organogenesis induction is a vital method to regenerate explants and produce complete organisms. In this study, we analyzed the applicability of three different root biostimulants and a commercially available synthetic rooting hormone (RH) for root organogenesis induction in *Epipremnum aureum* stem cuttings. The biostimulants used were *Aloe vera* gel (AV1), and garlic (GR2) and turmeric extracts (TM3), and the synthetic RH (TakeRoot®) used contained an active ingredient, indole butyric acid (0.01%). The *E. aureum* stem cuttings were placed in hydroponic pots and root development rates were monitored for up to 30 days. Recorded data from five parameters were analyzed: (1) number of rooted cuttings, (2) number of roots per stem cutting, (3) length of the longest and (4) shortest roots of the cuttings, and (5) rooting time. Stem cuttings were quantified using ImageJ software. The results showed that compared to the application of TakeRoot®, treatment with the biostimulant AV1 produced the longest roots, whereas stem cuttings treated with GR2 and TM3 did not produce significant results. Moreover, AV1 induced root organogenesis 16.67% faster than did TakeRoot® but no significant difference (*p*<0.05) was observed in the case of number of roots promoted per cutting. This study provides scientific evidence for the application of naturally derived RHs in the propagation of stem cuttings. Furthermore, *Aloe vera* gel, known for plant growth benefits, is the best choice for plant root propagation.

## Introduction

*De novo* organogenesis is one of the survival strategies of plants. This phenomenon occurs when adventitious shoots or roots are regenerated from detached or wounded plant tissues or organs (Xu & Huang, 2014; Sang et al., 2018). De novo shoot and root organogenesis can be induced in both tissue culture and the presence of natural elements. In the tissue culture techniques, the roots or shoots of explants are cultured in nutrient-enriched medium with suitable plant hormones (Skoog & Miller, 1957). In contrast, detached organs can naturally produce adventitious shoots and roots, which eventually grow into complete plant organisms. However, endogenous hormones are necessary for organogenesis (Ċosiċ et al., 2015; Hu et al., 2017). Examples of endogenous plant hormones include abscisic acid, auxin, cytokinin, gibberellic acid, and indole acetic acid.

Induction of adventitious roots is an essential step during plant propagation. Plant propagation is a technique for creating new plants from explants and can be performed using several methods. The reproduction or propagation of plants occurs either from seeds or vegetative means, such as cuttings (leaf, stem, or root), division layering, grafting, budding, and tissue cultures. Usually, rooting hormones (RHs) can be used as growth regulators by improving the overall rooting rate, production of adventitious roots, and the number and quality of roots during the propagation of plant stem cuttings (De Klerk, 2002). These growth regulators can be biological or artificial. Endogenous auxins are produced during plant organogenesis (Dubrovsky et al., 2008), while indole-3-butyric acid (IBA) and naphthalene acetic acid are the most frequently used synthetic growth hormones (Elmongy et al., 2018; Ludwig-Müller, 2000). Although synthetic RHs have been demonstrated to be effective, another method to propagate roots is by applying alternative RHs or biostimulants. In this study, the term ‘biostimulants’ refers to natural substances that can induce root organogenesis on stem cuttings. Studies have reported that natural ingredients, such as coconut water (Baque et al., 2011) and cinnamon (Kowalska et al., 2020), promote and enhance plant growth and are suitable substitutes for synthetic RH. Presently, many horticulturists use fresh *Aloe vera* gel to induce root growth in stem cuttings and air layering of plants (Fernando & Mirihagalla, 2021).

This study aimed to compare the effects of biostimulants and synthetic RH on the induction of root organogenesis in *Epipremnum aureum* stem cuttings as models. *Epipremnum aureum*, also known as the pothos plant, is a perennial plant with ornamental value. The plant belongs to the family Araceae and is known for its glossy, heart-shaped leaves with yellow or white variegation and its evergreen vine. The effects of biostimulants and synthetic RH were analyzed based on five different parameters: (1) number of rooted cuttings, (2) number of roots per stem cutting, (3) length of the longest and (4) shortest roots, and the (5) rooting time. In this study, the biostimulants used were *Aloe vera* gel, garlic and turmeric extracts, and a commercially available synthetic RH (TakeRoot®) containing IBA, 0.01% as the active ingredient was used. The results of this study provide an ideal alternative RH for use in the propagation of *E. aureum* plants, thus providing scientific recommendations to horticulturists for better cultivation of their explants.

## Materials and methods

### Preparation of Stem Cuttings

*Epipremnum aureum* plants were purchased online from a garden site in Cainta, Province of Rizal, Philippines. The plants were checked for suitability for experiment. A separate plant sample was submitted to Jose Vera Santos Memorial Herbarium (PUH) at the University of the Philippines for plant identification. A total of 75 stem cuttings were prepared in the laboratory. It was divided into 45 cuttings for the experimental groups (three subgroups) and 15 for each control group or triplicates with five stem cuttings per replication and labeled EA01 to EA05. To prepare the cuttings, a sterilized scalpel was used and the stem was cut to approximately five inches in length at a 45° angle with one node and leaf per stem. The cuttings were sprayed with Dithane M-45 fungicide to prevent fungal contamination of the plant tissues and were subjected to further analysis.

### Preparation of Biostimulants

In this study, *Aloe vera* gel (AV1) and garlic (GR2) and turmeric (TM3) extracts were used as the biostimulants. To remove dirt, the fresh, mature leaves of *Aloe vera* were thoroughly washed with sterile distilled water, and the thick epidermis was cut lengthwise into pieces. The gel was collected in a sterile container and ground using a blender to produce a watery mixture, and then transferred to a flask at a final concentration of 10 g/L. The garlic extract was prepared by grinding 10 g of peeled garlic cloves with 10 mL of water, followed by filtering to remove impurities and serial dilution to a final concentration of 50 g/L. For the turmeric extract, 10 g of crushed turmeric was diluted in 10 mL distilled water, and an extract of 25 g/L was prepared for further analysis.

### Propagation of Stem Cuttings and Data Analysis

Hydroponic pots were filled with 250 mL biostimulants. TakeRoot® (Garden Safe® brand) is ready-made hormone powder; therefore, no further preparation was required. Briefly, *E. aureum* stem cuttings were placed in hydroponic pots with ensuring that the cut ends were submerged. The cuttings were regularly misted with fungicides to prevent fungal contamination. Distilled water without biostimulant or synthetic RH was used as the blank control. The hydroponic pots were placed in a customized greenhouse that received sufficient light, but not direct sunlight. The day temperature and humidity were 32°C and 97%, respectively. The night temperature and humidity were 23°C and 42%, respectively. The parameters measured were: (1) number of rooted cuttings, (2) number of roots per stem cutting, (3) length of the longest root of cuttings, (4) length of the shortest root of cuttings, and (5) rooting time. Data recorded every 5 d. Cuttings with root lengths of ≥1 cm were considered rooted. The experimental setup was regularly monitored for up to 30 days, and the root growth was photographed using a digital camera. Root measurements were quantified using ImageJ (Image Processing and Analysis in Java) bundled with Java 1.8.0_172. Data were interpreted using ANOVA for a factorial complete randomized design, and the significance of results was confirmed using the Duncan’s multiple range test (DMRT).

## Results

In Figure 1 illustrates root organogenesis in stem cuttings of *E. aureum* after 30 d of the experiment under the influence of biostimulants and TakeRoot®, with the exception of biostimulant GR2. The concentration of GR2 was 50 g/L, and the pH ranged from 5.1 to 5.6. As a result, the cuttings did not endure the potentially lethal conditions within this pH range. Significant root formation was observed in stem cuttings treated with TakeRoot® and the other two biostimulants. Synthetic RH promoted root growth in 93.33% *E. aureum* stem cuttings, whereas biostimulants AV1 and TM3 induced roots in 86.67% and 33.33% stem cuttings, respectively (**Table 1**). TM3 induced the least amount of root organogenesis. Among the biostimulants, the AV1 at a concentration of 10 g/L produced an average of three roots per *E. aureum* stem cutting. TM3 at a concentration of 25 g/L produced one root/stem cutting. The stem cuttings treated with TakeRoot® initiated four roots/stem cutting (25% higher than the biostimulant AV1). Based on these findings, it appears that there is no significant difference (*p*<0.05) between biostimulant AV1 and synthetic RH in terms of the number of roots promoted in the stem cuttings.

**Fig. 1.**
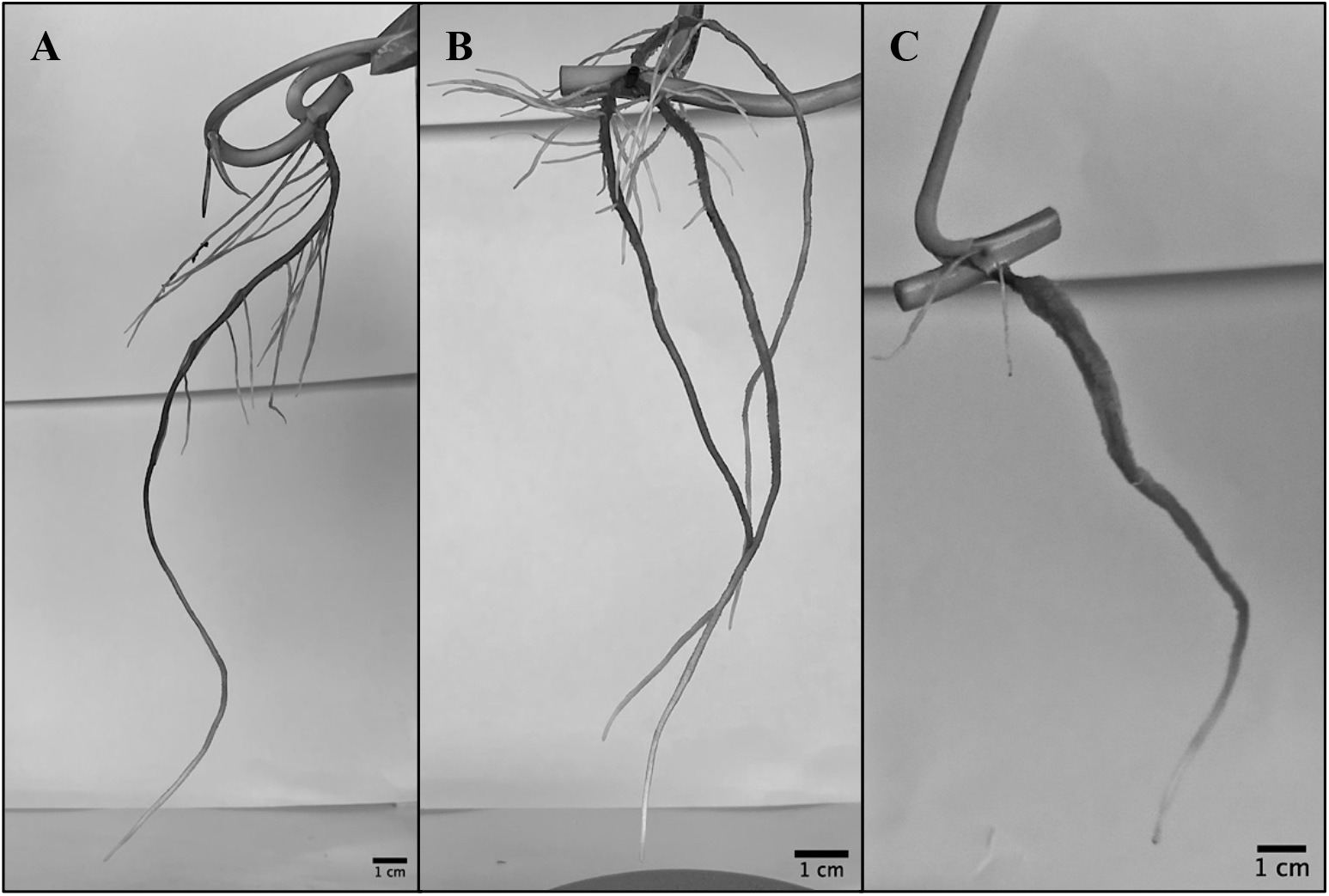
Roots of *E. aureum* stem cuttings after 30 d-experiment under the influence of (A) AV1, (B) TM3, and (C) TakeRoot® treatments. Longest root measurements were (A) 24.26 cm in EA01, (B) 12.05 cm in EA03, and (C) 11.78 cm in EA05.

**Table 1.**
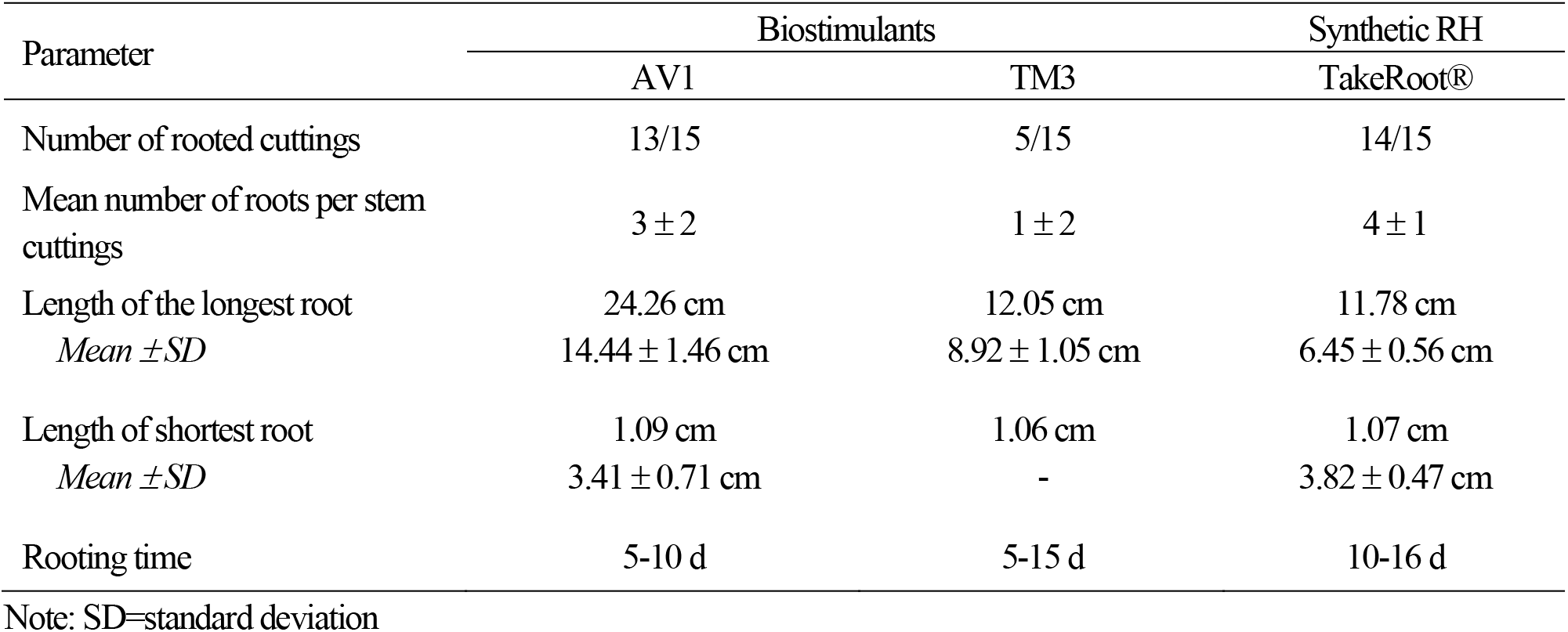
Summary of all root-related traits of the *E. aureum* stem cuttings (n=15) under biostimulants (AV1 and TM3) and TakeRoot® treatments.

| Parameter | Biostimulants |  | Synthetic RH |
| --- | --- | --- | --- |
|  | AV1 | TM3 | TakeRoot® |
| Number of rooted cuttings | 13/15 | 5/15 | 14/15 |
| Mean number of roots per stem cuttings | 3 ± 2 | 1 ± 2 | 4 ± 1 |
| Length of the longest root | 24.26 cm | 12.05 cm | 11.78 cm |
| Mean ± SD | 14.44 ± 1.46 cm | 8.92 ± 1.05 cm | 6.45 ± 0.56 cm |
| Length of shortest root | 1.09 cm | 1.06 cm | 1.07 cm |
| Mean ± SD | 3.41 ± 0.71 cm | - | 3.82 ± 0.47 cm |
| Rooting time | 5-10 d | 5-15 d | 10-16 d |

Note: SD=standard deviation

Both biostimulants and synthetic RH effectively increased root length (Table 2). ImageJ software was used to quantify adventitious root (AR) length. The longest recorded AR measurement was 24.26 cm for EA01 treated with AV1 (Fig. 1). Analyses showed that the mean root length in stem cuttings treated with AV1 (mean = 14.44 ± 1.46 cm) was significantly more (*p*<0.05) than that in cuttings treated with the synthetic RH (mean = 6.45 ±

**Table 2.**
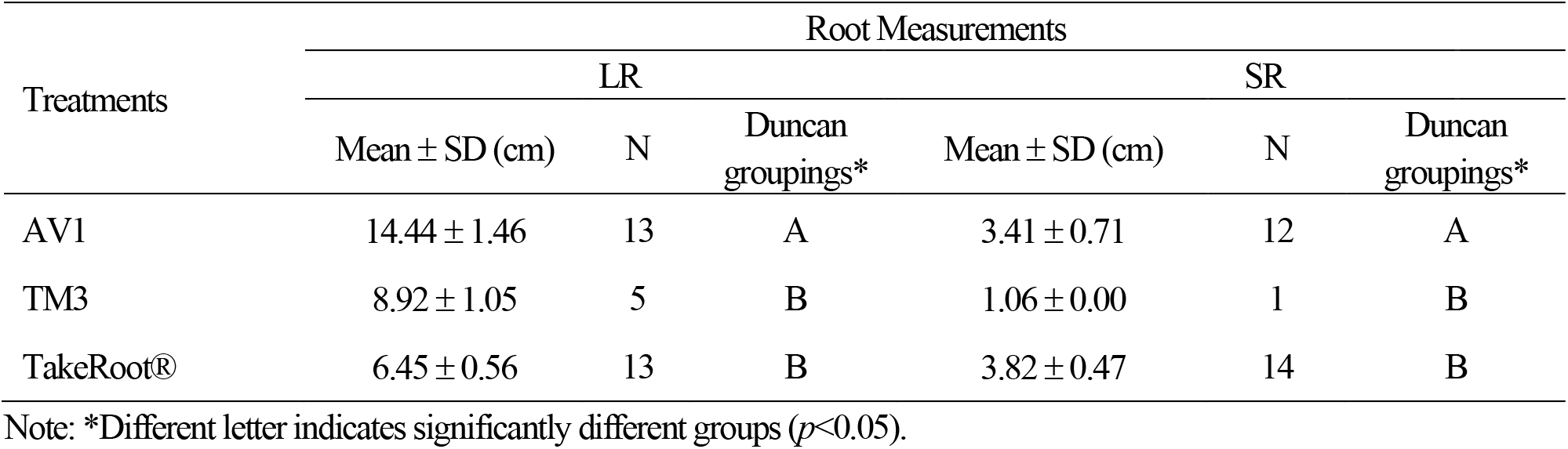
DMRT and mean values of the longest (LR) and shortest (SR) root measurements of *E. aureum* stem cuttings under biostimulants (AV1 and TM3) and TakeRoot® treatments after the 30-day experiment.

0.56 cm). This suggests that the *Aloe vera* gel stimulated root cell division in the elongation zone during organogenesis. *Aloe vera* gel extract contains vitamins, enzymes, amino acids, sugars, plant sterols, gibberellins, and salicylic acid. These nutrients are associated with improved vegetative growth, plant mineral composition, and oil production (Hamouda et al., 2012; Chatterjee et al., 2013; El Sheriff, 2017). In contrast, the shortest measurement of AR, spanning only 1.06 cm, was observed in EA05 treated with TM3 (Table 2). The biostimulant TM3 produced a short AR with a mean length of 0.08 cm. Furthermore, the Duncan’s test revealed that root measurements in stem cuttings treated with TM3 were significantly different (*p*<0.05) compared to those with AV1 and synthetic RH treatments. The root development rates of *E. aureum* stem cuttings with biostimulants and synthetic RH were monitored over time. The root measurements were recorded every 5 d for up to 30 d, as presented in Table 2. Stem cuttings treated with AV1 and TakeRoot® produced adventitious roots within 5-10 d, and 10-16 d, respectively. However, AV1 promoteed root organogenesis 16.67 % faster than the synthetic rooting hormone.

## Discussion

Root organogenesis induction in *E. aureum* stem cuttings with biostimulants, and synthetic RH was observed to be significant based on all five parameters: (1) number of rooted cuttings, (2) number of roots per stem cutting, (3) length of the longest and (4) shortest roots, and (5) rooting time. Among the biostimulants used in this study, the effects of *Aloe vera* gel was statistically comparable to those of TakeRoot® RH. The application of *Aloe vera* gel in plant propagation and cultivation has long been known in the field of horticulture. The gel contains vital elements to support plant growth. These include enzymes, carbohydrates, vitamins, amino acids, and plant hormones. Additionally, the gel exhibits antimicrobial activity against plant-pathogenic fungi (Hayat et al., 2016; Uda et al., 2018). Although the precise mechanisms of action of these plant extracts have not been fully elucidated, several of their constituents have been demonstrated to influence root growth or nutrient intake, both of which may affect the development of roots in the *E. aureum* stem cuttings. Sugars have a fundamental regulatory function in driving plant development. For instance, the plant target of rapamycin promotes growth in response to high sugar levels, while plant Snf1-related kinase 1 is particularly active upon sugar deprivation. Target of rapamycin and Snf1-related kinase 1 activities are modulated by the sugar status of a plant (Baena-González & Hanson, 2017; Rodriguez et al., 2019). Other constituents such as amino acids are coupled with increased nutrient uptake via chelation (Halpern et al., 2015; Souri & Hatamian, 2019). Although plant hormones have been identified in *Aloe vera* gel extracts, these are often in small and varying quantities (Surjushe et al., 2008; El Sherif, 2017; Wise et al., 2020). Auxin is a primary growth-promoting hormone that is involved in AR initiation. An example is indole-3-acetic acid, an auxin produced during plant cellular mechanisms, which promotes AR formation and auxin homeostasis through the rooting process (Druege et al., 2016; Gonin et al., 2019). Therefore, when applied to *E. aureum* stem cuttings, the AV1 imparts root stimulating actions either indirectly through stress mediation and chelation or directly through hormones and carbon supply.

Based on the literature, biostimulants AV1 and TM3 are enriched in minerals and trace elements necessary for root growth and development. The analytical result showed that *Aloe vera* belongs to a group of calcium-rich vegetables, which contains slightly increased levels of Na, P, and Fe (Yamaguchi et al., 1993). Moreover, other elements such as Al, B, Mg, Mn, S, and Sr were also detected (Yamaguchi et al., 1993). The mineral elements present in turmeric (*Curcuma longa* L.) are K, Ca, P, Mg, and Na (Jabborova et al., 2021). Together with N, P and K have been observed to enhance shoot and root growth of wheat, upland rice, and corn (Fageria & Moreira, 2011). However, although P stimulates the early root growth rate, Fe or Zn reduces it (Bouain et al., 2018). Nevertheless, root organogenesis in *E. aureum* stem cuttings was induced by biostimulants, particularly AV1, which showed significant results for all parameters. Further analysis is recommended to determine the effects of turmeric extract on plant growth and development.

The synthetic RH TakeRoot® is a commercially available RH powder containing the active ingredient IBA (0.01%). The effect of AV1 was found to be statistically comparable to that of TakeRoot® for all parameters, except rooting time. Synthetic RH is a chemically synthesized artificial product that resembles the natural hormones of a plant and stimulates root growth. IBA is the common synthetic compound in many commercially available horticultural rooting products that elicits auxin-like effects, such as root initiation, stem bending, and leaf epinasty (Frick & Strader, 2018). Compared to other studies, in this study, the concentration of IBA was low, only 0.01%. A previous study found that IBA at a level of 20% showed the best results in terms of the number of leaves per plant, the number of roots per plant, root diameter, and survival percentage in *Clerodendrum splendens* (glory tree) stem cuttings (Jamal et al., 2016). Likewise, a 0.2% IBA gel strongly influenced the success and quality of adventitious rooting in cannabis cuttings, delivering a higher rooting success rate than with the 0.2% willow extract (Caplan et al., 2018). Thus, the exogenous application of IBA in TakeRoot® enhances root organogenesis in *E. aureum* stem cuttings in terms of the number of rooted cuttings, roots per cutting, and root length.

In conclusion, the *Aloe vera* gel extract at a concentration of 10 g/L and synthetic RH TakeRoot® with IBA, 0.01% effectively induced root organogenesis in *E. aureum* stem cuttings. Both treatments produced a substantial number of roots (3-4 roots per cutting) and the development of roots with high success rates (AV1 = 86.67%; TakeRoot® = 93.33%). However, root formation in cuttings treated with TakeRoot® is slower than that in cuttings treated with AV1. Further study on the possible application of turmeric extract at much higher concentrations or as additive with the existing rooting hormone is recommended because TM3 contains minerals and nutrients necessary for plant growth and development. This study provides scientific evidence on the application of naturally derived RHs, such as *Aloe vera* gel extract, to induce root organogenesis in plant cuttings.

## Data availability

The authors confirm that all relevant data supporting the findings of this study are available in Mendeley Data with DOI 10.17632/bdvyz262fb.1

## Acknowledgements

The authors would like to thank Ms. Trisha Gotinga of Research and Development Innovation Center, Our Lady of Fatima University-Quezon City Campus (RDIC-OLFU), and Mr. Mark Stephen Real, Dr. Rose Marie Santos, and Mrs. Bernardita Gacutan of Department of Biology for providing technical support on experimental planning, writing and data analysis.

## Author contribution

D.E.V. designed research, performed experiments, analyzed data, and wrote the manuscript. E.A., A.B., R.D., N.L.M., M.A.S. performed experiments and helped writing the manuscript. R.D.J. designed and supervised the research, analyzed data, and wrote the manuscript. All authors approved the final version of the manuscript.

## References

Baena-González, E. and J. Hanson, 2017. Shaping plant development through the SnRK1–TOR metabolic regulators. Curr. Opin. Plant Biol., 35: 152–157.

Baque, A., Y.K. Shin, T. Elshmari, E.J. Lee and K.Y. Paek, 2011. Effect of light quality, sucrose and coconut water concentration on the microporpagation of calanthe hybrids (“bukduseong” × “hyesung” and “chunkwang” × ‘hyesung’). Aust. J. Crop Sci., 5: 1247–1254.

Bouain, N., A. Korte, S.B. Satbhai, S.Y. Rhee, W. Busch and H. Rouached, 2018. Systems approaches provide new insights into Arabidopsis thaliana root growth under mineral nutrient limitation. BioRxiv, 460360.

Caplan, D., J. Stemeroff, M. Dixon and Y. Zheng, 2018. Vegetative propagation of cannabis by stem cuttings: effects of leaf number, cutting position, rooting hormone, and leaf tip removal. Can. J. Plant Sci., 98: 1126–1132.

Chatterjee, P., B. Chakraborty and S. Nandy, 2013. Aloe vera: review with significant pharmacological activities. Mintage J. Pharm. Med. Sci., 1: 21–24

Ċosiċ, T., V. Motyka, M. Raspor, J. Savić, A. Cingel, B. Vinterhalter, D. Vinterhalter, A. Trávníčková, P.I. Dobrev, B. Bohanec and S. Ninković, 2015. In vitro shoot organogenesis and comparative analysis of endogenous phytohormones in kohlrabi (Brassica oleracea var. gongylodes): effects of genotype, explant type and applied cytokinins. Plant Cell Tissue Organ Cult., 121: 741–760.

De Klerk, G.J. 2002. Rooting of microcuttings: theory and practice. In Vitro Cell Dev. Biol. Plant., 38:415–422.

Druege, U., P. Franken and M.R. Hajirezaei, 2016. Plant hormone homeostasis, signaling, and function during adventitious root formation in cuttings. Front. Plant. Sci., 7: 1–14.

Dubrovsky, J.G., M. Sauer, S. Napsucialy-Mendivil, M.G. Ivanchenko, J. Friml, S. Shishkova, J. Celenza and E. Benková, 2008. Auxin acts as a local morphogenetic trigger to specify lateral root founder cells. Proc. Natl. Acad. Sci. USA., 105: 8790–8794.

El Sherif, F. 2017. Aloe vera leaf extract as a potential growth enhancer for populus trees grown under in vitro conditions. Am. J. Plant Biol., 2: 101–105.

Elmongy, M.S., Y. Cao, H. Zhou and Y. Xia, 2018. Root development enhanced by using indole-3-butyric acid and naphthalene acetic acid and associated biochemical changes of in vitro azalea microshoots. J. Plant Growth Regul., 37: 813–825.

Fageria, N.K. and A. Moreira, 2011. The role of mineral nutrition on root growth of crop plants. Adv. Agron., 110: 251–331.

Fernando, M. and M. Mirihagalla, 2021. Effect of Aloe vera gel for inducing rooting of stem cuttings and air layering. J. Dry Zone Agric., 1: 13–26.

Frick, E.M. and L.C. Strader, 2018. Roles for IBA-derived auxin in plant development. J. Exp. Bot., 69:169–177.

Gonin, M., V. Bergougnoux, T.D. Nguyen and P. Gantet, 2019. What makes adventitious roots? Plants, 8:1–24.

Halpern, M., A. Bar-Tal, M. Ofek, D. Minz, T. Muller and U. Yermiyahu, 2015. The use of biostimulants for enhancing nutrient uptake. Adv. Agron., 130: 141–174.

Hamouda, A.M.A., D.M.G. Hendi and O.F. Abu El-Leel, 2012. Improving basil growth, yield and oil production by Aloe vera extract and active dry yeast. Egypt J. Hort., 39: 45–71.

Hayat, M.K., T. Kumar, N. Ahmad, S.A. Jan, W. Sajjad, S. Faisal, B.X. Gao, K. Khan, Z.U. Rahman and B.H. Abbasi, 2016. In vitro antimicrobial activity of Aloe vera L. extracts against pathogenic bacteria and fungi. Mycopath, 14: 21–27.

Hu, W., S. Fagundez, L. Katin-Grazzini, Y. Li, W. Li, Y. Chen, X. Wang, Z. Deng, S. Xie, R.J. McAvoy and Y. Li, 2017. Endogenous auxin and its manipulation influence in vitro shoot organogenesis of citrus epicotyl explants. Hortic. Res., 4.

Jabborova, D., R. Choudhary, R. Karunakaran, S. Ercisli, J. Ahlawat, K. Sulaymanov, A. Azimov and Z. Jabbarov, 2021. The chemical element composition of turmeric grown in soil–climate conditions of Tashkent region, Uzbekistan. Plants, 10: 1–11.

Jamal, A., G. Ayub, A. Rhaman, A. Rashid, J. Ali and M. Shabab, 2016. Effect of IBA (Indole Butyric Acid) levels on the growth and rooting of different cutting types of Clerodendrum splendens. Pure Appl. Biol., 5: 64–71.

Kowalska, J., J. Tyburski, J. Krzymińska and M. Jakubowska, 2020. Cinnamon powder: an in vitro and in vivo evaluation of antifungal and plant growth promoting activity. Eur. J. Plant Pathol., 156: 237–243.

Ludwig-Müller, J. 2000. Indole-3-butyric acid in plant growth and development. Plant Growth Regul., 32: 219–230.

Rodriguez, M., R. Parola, S. Andreola, C. Pereyra and G. Martínez-Noël, 2019. TOR and SnRK1 signaling pathways in plant response to abiotic stresses: do they always act according to the “yin-yang” model? Plant Sci., 288.

Sang, Y.L., Z.J. Cheng and X.S. Zhang, 2018. Plant stem cells and de novo organogenesis. New Phytol., 218: 1334–1339.

Skoog, F. and C. Miller, 1957. Chemical regulation of growth and organ formation in plant tissues cultured in vitro. Symp. Soc. Exp. Biol., 11: 118–130.

Souri, M.K. and M. Hatamian, 2019. Aminochelates in plant nutrition: a review. J. Plant Nutr., 42: 67–78.

Surjushe, A., R. Vasani and D.G. Saple, 2008. Aloe vera: a short review. Indian J. Dermatol., 53: 163–166.

Uda, M.N.A., N. Harzana Shaari, N. Shamierasaid, N.H. Ibrahim, M.A.M. Akhir, M.K. Rabani Hashim, M.N. Salimi, M.A. Nuradibah, U. Hashim and S. Gopinath, 2018. Antimicrobial activity of plant extracts from Aloe vera, Citrus hystrix, Sabah Snake Grass and Zingiber officinale against Pyricularia oryzae that causes rice blast disease in paddy plants. IOP Conf. Ser. Mater. Sci. Eng., 318.

Wise, K., H. Gill and J. Selby-Pham, 2020. Willow bark extract and the biostimulant complex Root Nectar® increase propagation efficiency in chrysanthemum and lavender cuttings. Sci. Hortic., 263.

Xu, L. and H. Huang, 2014. Genetic and epigenetic controls of plant regeneration. Curr. Top. Dev. Biol., 108: 1–33.

Yamaguchi, I., N. Mega and H. Sanada, 1993. Components of the gel of Aloe vera (L.) Burm. f. Biosci. Biotechnol. Biochem., 57: 1350–1352.

